# A faster extrinsic incubation period could explain Usutu leading West Nile in temperate Europe

**DOI:** 10.64898/2026.04.02.716093

**Authors:** Robert S. Paton, Maisie Vollans, Lorna Glenn, Martyn Fyles, Alexander G.C. Vaux, Jolyon M. Medlock, Julie Day, Thomas Ward

**Author notes:** Co-first authors (Robert S. Paton, Maisie Vollans).

## Abstract

Usutu virus (USUV) is a mosquito-borne flavivirus that has recently expanded northwards in Europe and become endemic in the UK. USUV emergence often precedes the closely related West Nile virus (WNV), potentially reflecting differences in epidemiological parameters. One key parameter is the extrinsic incubation period (EIP), the time taken for a mosquito to become infectious after a blood meal. Here we present the first quantitative estimate of the temperature dependent EIP for USUV in the vector *Culex pipiens molestus*. We were able to quantify the shortening of the USUV EIP from 68.06 days (95% CrI: 38.62 to 141.04) at 17 °C to 12.35 days (95% CrI: 8.10 to 17.09) at 25°C. This was achieved by re-analysing laboratory data with a bespoke Bayesian model that accounted for important features of the experimental design. Under UK summer temperatures, the median EIP of USUV is 37.75% (95% CrI: 3.38 to 55.68) shorter than that of WNV, and the potential transmission season of USUV is both longer and geographically more extensive. Under RCP8.5 climate projections, WNV transmission suitability could match or exceed current USUV levels between 2055 and 2065, highlighting the future threat to the UK from emerging mosquito borne pathogens. Our findings support USUV as a precursor for WNV in Europe and provide a robust characterisation of a key epidemiological parameter of USUV, enabling accurate modelling of its transmission dynamics.

## Introduction

Usutu virus (USUV) is an arbovirus that primarily infects birds, causing high mortality and mass die-offs in species such as the Eurasian blackbird, *Turdus merula* [1]. The virus is maintained in an enzootic transmission cycle involving avian hosts and ornithophilic mosquitoes, with birds acting as amplifying hosts and reservoirs of the virus [2]. Spillover events can occur to humans and other mammals, however they are dead-end hosts and therefore do not contribute to onward transmission [3]. The dominant vector of USUV in Europe are mosquitoes from the *Culex pipiens* species complex [4], with *Culex pipiens molestus* biotype considered a potential bridge vector that facilitates the spillover of USUV from birds to humans in Europe, due to its more diverse biting habits including both birds and mammals [5].

Clinical symptoms of USUV infection are not well characterised in humans, due to the relatively low number of reported cases in comparison to other arboviruses, such as West Nile virus (WNV) [6, 7]. In the cases that have been reported, USUV infections are generally asymptomatic or result in mild symptoms. However, the virus can in rare cases cause severe neurological symptoms (including meningitis and meningoencephalitis) which are more likely to occur in immunocompromised individuals [3]. As a consequence, USUV is a notable and growing public health concern.

In recent years, USUV has undergone substantial range expansion in Europe. USUV is thought to have been introduced to Europe from Africa multiple times since the 1950s via the movement of infectious migratory birds [8]. The first documented case of USUV in birds in Europe dates from 1996 [9], where retrospective analyses of dead Eurasian Blackbirds (*Turdus merula*) were tested in Italy. Since its introduction, USUV has become widespread across Europe. This includes Northern Europe, where the United Kingdom first detected USUV in both birds and mosquitoes in 2020 [10, 11]. There is evidence to suggest that USUV overwinters in the UK, with autochthonous transmission among birds and vertical transmission to mosquito eggs in the *Culex pipiens* complex [12]. To date (early 2026), there have been no published reports of autochthonous human infections in the UK.

WNV was retrospectively detected in *Aedes vexans* sampled from Gamston, Nottinghamshire, UK from samples collected in 2023 [13], but with no subsequent reports of autochthonous transmission to either birds, or humans. This reflects broader trends observed across Central and Northern Europe, where USUV is often either transiently detected or becomes established endemically in avian populations several years before the first detection of WNV in birds or humans. This has been reported in several countries such as Austria [14, 15], Germany [16, 17] and the Netherlands [18, 19]. Similarly, USUV has been shown to overwinter and become endemic in multiple European countries, including Belgium and the UK, before autochthonous transmission of WNV was reported [12, 20].

Both USUV and WNV share hosts, vectors and importation routes. This suggests that the repeated observation that USUV circulation precedes WNV [4, 21] emergence may indicate differences in their transmission dynamics. However, there has been little quantitative work to test this hypothesis. A major barrier to comparative analysis between USUV and WNV is the poor characterisation of key epidemiological parameters for USUV. A key parameter that is yet to be characterised for USUV is the extrinsic incubation period (EIP), which is the time taken for a mosquito to become infectious after ingesting an infected blood meal.

During the EIP, the virus must overcome multiple tissue barriers within the vector, evade immune responses and disseminate to the mosquito salivary glands [22–24]. Empirically, the EIP is generally defined as the time from ingestion of an infectious blood meal to viral dissemination to either the mosquito’s legs or salivary glands. The detection of viral material in the mosquito body is typically used as evidence of infection only, not dissemination. Detection of virus in the legs is often preferred for determining the EIP, as it provides a more reliable proxy for the feasibility of transmission via mosquito saliva, given the technical challenges associated with saliva extraction [25].

The EIP is highly temperature sensitive: higher temperatures lead to faster rates of viral replication within the mosquito, shortening the time required for dissemination to the salivary glands [26–28]. As mosquitoes are poikilotherms, their internal body temperature is largely determined by ambient environmental conditions, thus external temperature strongly influences how rapidly mosquitoes become infectious. The EIP is not deterministic and there is substantial variation among individual mosquitoes, making it important to statistically characterise individual heterogeneity. The specific relationship between EIP and temperature has been characterised for multiple arboviruses and differs among them [26–28].

Here, we reanalyse experimental data from three previously published studies [5, 29, 30] in which adult *Cx. pipiens molestus* mosquitoes were experimentally infected with live USUV and maintained under different, constant temperature conditions. We develop a bespoke Bayesian model to estimate the temperature dependent extrinsic incubation period for USUV. This framework mainly relies on body positivity data, applying a modifier to account for the delay between body infection and dissemination. We use this model, alongside a previously published model of the temperature-dependent West Nile virus EIP [28], to compare the feasibility of transmission of each virus in the UK. This allows us to examine whether differences in the EIP may help explain the establishment of USUV. For both USUV and WNV, we evaluate a minimum condition for transmission: the probability the EIP could complete in the average mosquito lifespan, giving opportunity for onward infection. We combine these analyses with climate projections to explore how the relative success of WNV and USUV may change under a high-emissions scenario, RCP 8.5 [31]. To our knowledge, this work provides the first characterisation of the temperature-dependent EIP of USUV and the first quantitative comparison of this key epidemiological parameter with WNV.

## Methods

### Laboratory protocols

This analysis used data from three laboratory studies: Pilgrim *et al.* (2024) [5], Seechurn *et al.* (2025) [29] and Holicki *et al.* (2020) [30].

Pilgrim *et al.* (2024) [5] and Seechurn *et al.* (2025) [29] experimentally infected cohorts of *Culex pipiens molestus*with the UK-endemic strain of USUV (London, 2020) via infectious blood meals. Pilgrim *et al.* (2024) [5] infected a total of 187 mosquitoes, while Seechurn *et al.* (2025) [29] infected 336 mosquitoes. Following infection, mosquitoes were divided into groups, and held at multiple different, constant temperatures representative of the UK climate (17°C to 23°C). At predefined days post-infection (dpi), subsets of mosquitoes were sampled, and bodies and saliva tested for viral RNA using quantitative reverse transcriptase PCR (RT-qPCR).

A limitation of both studies is that the saliva positivity shows no clear trend through time, or across temperatures. This observation is consistent with a previous study which demonstrated that forced salivation assays have technical challenges and poor reproducibility, often leading to an underestimation of transmission compared with using viral dissemination to mosquito legs as a proxy [25]. For these reasons, our previously published model describing the EIP of WNV used leg positivity as a proxy for disseminated infection as opposed to saliva [28]. In this study we therefore do not model the saliva data from Pilgrim *et al.* (2024) [5] or Seechurn *et al.* (2025) [29], instead supplementing the body positivity data with another source.

Holicki *et al.* (2020) [30] followed a similar experimental procedure to Pilgrim *et al.* (2024) [5] and Seechurn *et al.* (2025) [29] but for only one temperature treatment (25°C). A total of 42 *Cx. pipiens molestus* from German and Serbian colonies were infected with USUV Africa 2 strain and tested for viral RNA using RT-qPCR. A crucial difference however is that this study analysed viral RNA separately in mosquito bodies and the legs / wings. This study therefore contributed an additional 25°C condition to model the process of whole-body infection, alongside data we could use to parametrise disseminated infection in the legs and wings, which we explain later in the methods.

### Data treatment and interpretation

PCR assays amplify viral RNA, irrespective of whether it is live, infectious virus or inert viral material. This contrasts with other vector competence methods, such as plaque assays, which assess whether infectious virus can be cultured from a sample. For example, in Kilpatrick *et al.* (2008) [32] *Culex pipiens pipiens* mosquitoes were infected with WNV and plaque assays were used to test whole body, leg and saliva at multiple dpi. In our published reanalysis [28] of these plaque assay data, we modelled the temperature-dependent EIP of WNV by treating positive tests as interval censored observations (with dissemination occurring sometime between ingestion of an infectious bloodmeal and sampling) and negative tests as right censored (dissemination having not yet occurred) observations. Our model assumed that positivity of disseminated infection is low soon after infection, when tests often precede dissemination, and increases over time as the virus disseminates throughout the mosquito. We found that positivity increases more quickly at higher temperatures as viral replication is faster. We make similar assumptions in this work.

Estimating the temperature-dependent EIP for USUV presents additional challenges, as available data are from RT-qPCR outputs rather than plaque assay. First, the body positivity data show a complex pattern of positivity following infection, for details please see **Figure S1**. Body positivity is initially 100% (demonstrated by Seechurn *et al.* (2025) [29], who sampled mosquitoes on day zero post infection) followed by a decline soon after, before subsequently increasing. This trend could be explained by an artefact of the experimental design: initial RT-qPCR positivity reflects viral RNA from the bloodmeal that has not yet infected the mosquito. As the bloodmeal is metabolised, it is expected this source of viral RNA decreases, resulting in the observed decline in positivity. Subsequent increases in positivity reflect the virus infecting the midgut and replicating: this infection process is what we aim to uncover. This motivates an interesting modelling problem, whereby positivity is driven by both bloodmeal derived RNA and true infection, changing in representation through time.

Second, the body positivity data from Seechurn *et al.* (2025) [29] shows an unexpected trend with temperature. Mosquitoes held at 19 and 22°C show higher positivity across all dpi sampling points than those held at 20°C. As viruses typically replicate faster as temperatures increase, the observed dip in positivity at 20°C is unlikely to be genuine signal and we therefore assume that the 20°C experimental treatment by in large failed. However, RNA would still be returned as positive in early tests due to residual viral RNA from the blood meal, even in the absence of a viable infection. We therefore treat the data from the 20°C infection treatment in Seechurn *et al.* (2025) [29] as reflecting exclusively bloodmeal derived positivity, with no infections possible. This helps us to parameterise the decay in positivity associated with viral RNA only.

To address these challenges, we decided to model the process of mosquito whole body infection using six temperature treatments: 17, 21, 23°C from Pilgrim et al. (2024) [5] in combination with the 18 and 22°C temperature from Seechurn *et al.* (2025) [29] and 25°C treatment from Holicki *et al.* (2020) [30]. These data were available from the original publications and were extracted for each dpi and temperature condition. Then we used the leg and wing data available from Holicki *et al.* (2020) [30] to model the process of the mosquito developing a disseminated infection, by estimating a modifier to the body infection process to reflect the delay in the infection becoming systemic.

### Model overview

Each observed mosquito body is tested on a particular day (*d*_*i*_ where *i* denotes the unique individual mosquito) and receives either a positive 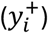 or negative 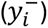 RT-qPCR test result. The test result determines whether the mosquito became infected between the start of the study and the day it was observed (positive when observed; interval censored) or may become infected in the future (negative when observed; right censored). However, positive results also reflect residual viral RNA in the blood meal, while negative results may include mosquitoes that never develop infection, even if given an infinite time horizon.

#### Bloodmeal positive tests

To account for residual viral RNA from the blood meal, we define an exponential decline in the probability of a positive test resulting from the blood meal, starting the day after ingestion:

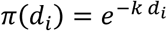

The strictly positive parameter *k* describes the rate of decline from 100% bloodmeal positive rate on day zero (because infections will not begin immediately) with π(d_i_) the probability of a bloodmeal positive result on day *d*_*i*_. We chose this functional form to reflect changes in RNA concentration through time in a parsimonious way [33, 34]

#### Incomplete infection rates

Not all mosquitoes will necessarily develop an infection: variation in mosquito immunity, microbiome and the viral load in the blood meal may mean that, even with unlimited time, some mosquitoes may remain uninfected. This is relevant for right censored observations, where it is unknown whether an infection will ever develop. We therefore estimate a study specific parameter, ρ_*s*_, which gives the probability that a mosquito would ever become infected (irrespective of time). This also allows the model to reflect differences in the two experimental protocols in delivering an infectious dose to sampled mosquitoes.

#### Model structure

In our model we use a Weibull distribution (parametrised by a shape parameter *⍺* and scale parameter *s*) to describe the variation in the approximate EIP across individual mosquitoes. We chose this specific distribution as our recent work supported this as the best descriptor of the West Nile virus extrinsic incubation period in *Culex pipiens* [28].

We assume that the median time to infection μ*T*^′^ varies log-linearly with temperature. We standardise experimental temperature by the mean (*T̅*) and standard deviation (*sd*(*T*)) to improve convergence properties in the model:

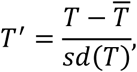

and the log-median is modelled as:

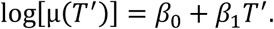

For a Weibull distribution with shape *⍺* and scale ϕ(*T*^′^), the median satisfies μ*T*^′^= ϕ(*T*^′^)(log(2))^1/*⍺*^. Therefore, the scale parameter is:

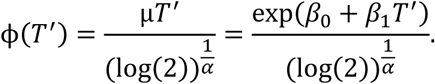

We model the process of positives due to residual viral RNA from the bloodmeal using data from the 20°C temperature treatment in Seechurn *et al.* (2025) [29], where infection rates were very low:

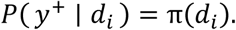

Mosquito bodies testing positive in other temperature treatments are a mixture of ‘true positives’ (mosquitoes that have developed infection) and ‘bloodmeal positives’ (residual RNA detected from the blood meal). First, we define *F*[ *d*_*i*_ ∣ *⍺*, *ϕ*(*T*^′^) ] as the cumulative probability of the Weibull distribution, modelling the interval censored observation of a true positive (interval censoring occurs as an infection could have occurred at any point between ingestion of the infectious blood meal, and the time of the positive test result). Then we define bloodmeal positive tests as *A* (where the probability is 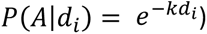 and true positives as *B* (where the probability is given by 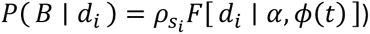.

As *A* and *B* are not mutually exclusive (*A* ∪ *B*) we model positive tests as:

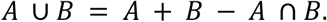

We assume independence between residual blood-meal RNA detectability (event *A*) and the occurrence of true infection by day *d* (event *B*), so that *P*(*A* ∩ *B*) = *P*(*A*|*d*) *P*(*B*|*d*).

Under the assumption that *A* and *B* are independent, *P*(*A* ∩ *B*) = *P*(*A*)*P*(*B*). This allows us to write the probability of an observed positive test on day *d*_*i*_ in study *s*_*i*_ in temperature treatment 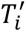 as:

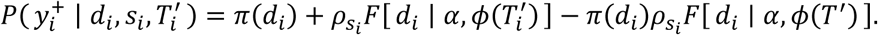

Here, 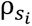 was modelled as a random effect, with each level corresponding to one of the three studies:

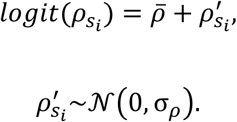

#### Wing and leg positivity

We assume that being body positive is a prerequisite for testing positive in the wings / legs, and that disseminated infections cannot be a result of residual viral RNA from the bloodmeal. Wing and leg positive tests 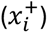 are therefore the outcome of the overall infection rate 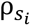 and a cumulative probability of infection. As we have no temperature dependent wing / leg positivity data, we rely on a modified body infection process to describe this cumulative probability:

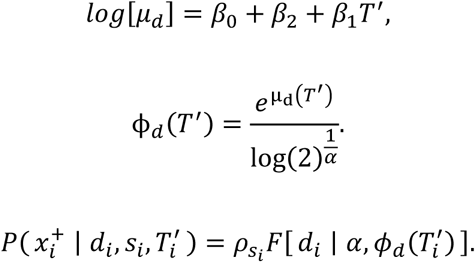

Here the median μ_d_ is a modified version of μ with an additional parameter β_2_ which allows the model to correct the median response for body infections to account for the further delay associated with disseminated infection. This adjustment was applied to the median response, so relies on the same scale parameter *⍺* as the body infection process.

#### Likelihood

To capture additional variation associated with test outcomes, we modelled the likelihood (ℒ) as a Beta-binomial process. Here, the mean probability *p* and dispersion parameter θ give the two positive shape parameters (*s*_1_ and *s*_2_) of a Beta distribution allowing additional variation in outcomes across the number of positive tests (n) out of the total tests (*N*):

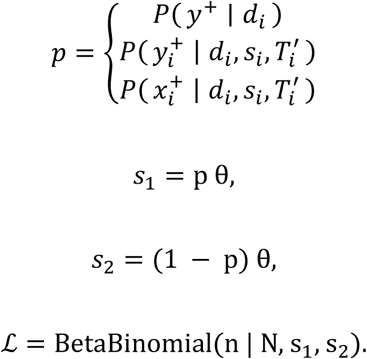

#### Model selection

We compare the model structure described above to alternative formulations that reflect different hypotheses. First, we developed a model where the approximate EIP_50_ did not change with temperature, such that:

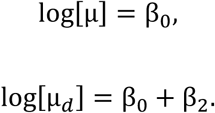

Second, we assume that the approximate EIP_50_ was fixed across temperatures, but the probability of ever becoming infected is a function of temperature:

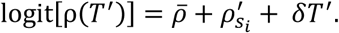

Here, a logit-linear function temperature dependent process (*δT*^′^) modifies the study specific probability of infection. This structure allows the model to explain differences in temperature treatments using the probability of infection only.

A final (and most complex model) allows the approximate EIP_50_ and *ρ* to vary with temperature. Models were compared using the leave-one-out cross validation information criterion, “looic” [35]. Models with looic scores within two standard errors were considered equivalent.

All models were written in the Bayesian programming language “stan” [36] and fit using Hamiltonian Monte Carlo methods interfaced through the statistical programming language “R” [37]. Priors used in the model are given in the **Supplementary Table 1**.

### Comparison to WNV

To determine how the transmission season for USUV compared to WNV in the UK, we compared the approximate EIP estimated in this paper to our previously published temperature dependent function for the median EIP of WNV [28]. We extracted hourly temperature values from ERA5-Land [38] using Google Earth Engine [39] for the years 2019 to 2024. We created a 5km-by-5km daily raster by resampling the mean of native-resolution pixels overlapping each output grid and averaging hourly values. For each pathogen, we solved the log-linear function describing the median EIP on daily temperatures for each year. We then calculated the geometric mean of these EIPs for each month in each pixel across the five years.

#### Survival models

Our study analyses the EIP of USUV in *Cx. pipiens molestus* while the model for the EIP of WNV used *Cx. pipiens pipiens* [28]. However, the values of EIPs in themselves are difficult to interpret in isolation, particularly when comparing across different vectors and pathogens. To accurately represent the dissemination process for both viruses we use a bespoke vector survival model for each vector / virus pair. This is challenging, as a component of the differences in the probability metric we calculate will be due to the vector, rather than the virus. We felt, however, that it was preferable to accurately represent the vector pathogen pair, as opposed to using the same survival model that would unavoidably be incorrect for one EIP model.

We used two experimental studies to estimate the temperature dependent survival of *Cx. pipiens molestus* and *Cx. pipiens pipiens* [40, 41]. In the first, Andreadis *et al.* (2019) [41] reared cohorts of *Cx. pipiens pipiens* under different temperature conditions (15, 20, 25, 27.5 and 30°C) with 20 males and 20 females in each temperature condition. We digitised data for each temperature treatment from Figure 1c for females only, allowing us to extract pseudo row-level data. A second study by Spanoudis *et al.* (2018) [40] examined more temperature conditions for *Cx. pipiens molestus* (15, 17.5, 20, 22.5, 25, 27.5, and 30) but we could only extract mean and standard error from Figure 4 in the main text. A full description of the temperature dependent survival models we developed for each species can be found in the Electronic Supplementary Materials.

**Figure 1:**
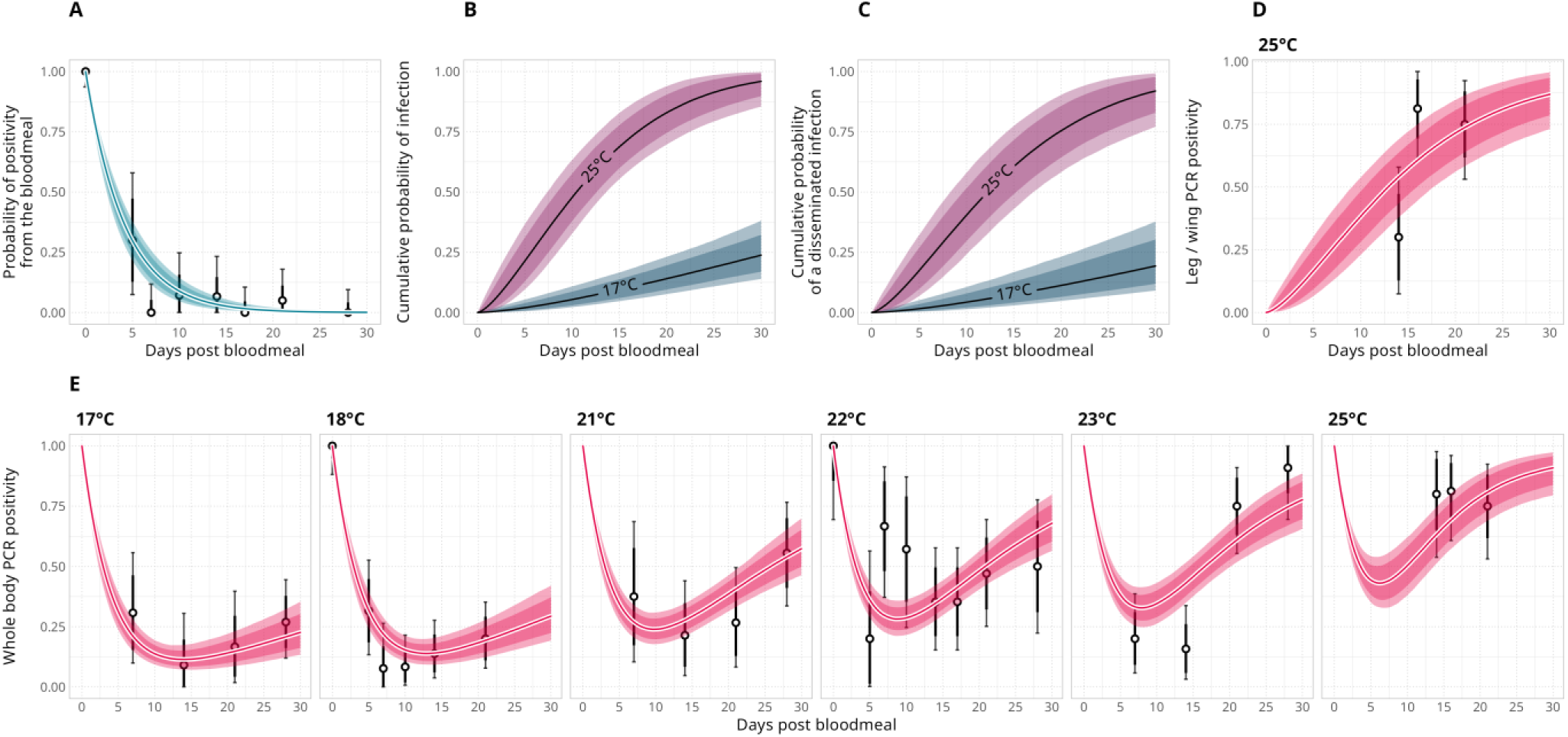
Model fit to data. *Panel A* shows the model estimated probability of a PCR positive test resulting from viral RNA in the bloodmeal. Points and intervals show test data from “day 0” post infectious bloodmeal (which are all exclusively bloodmeal positive) and the 20°C treatment from Seechurn *et al.* (2025), where we it appeared tests reflected almost exclusively bloodmeal positivity. *Panel B* shows the Weibull cumulative density function for the approximate EIP for USUV solved at the temperature extremes from the laboratory data. In *panel C* we show the cumulative probability corrected for disseminated infection. *Panel D*shows the model fit to the 25°C leg / wing positivity data from Holicki *et al.* (2020). Finally, *panel E* shows the model fit to the whole-body positivity data in each experimental treatment, where black points show the raw proportions and black lines are the credible intervals on these proportions. The model fit in *panel C* reflects the combination of the bloodmeal positivity function from *panel A*, the temperature dependent cumulative probability function in *panel B* and the study specific probability of infection ρ_*s*_.

#### Transmission risk

We use these survival models to determine the predicted average lifespan of each mosquito biotype and estimate the probability of the EIP completing within the vector’s lifespan across space and time. To estimate this, we calculated the average mosquito lifespan at the given temperature and used the corresponding temperature dependent pathogen cumulative probability function to estimate the probability of the EIP completing in that lifespan. We term the number of days where there is more than a 50% chance of competing within the average vector lifespan as “days of transmission risk”.

#### Forecasting the impact of climate change

We used the Met Office projections of RCP8.5 [31] for each year in decadal increments between 2025 and 2075 to assess how the transmission risk increases under climate change. Daily temperatures are only produced for RCP8.5, restricting our selection to this scenario. Projections are produced for each day across 12 model runs at a 12km x 12km resolution. We calculated the number of days across each year and model run where there was a 50% chance or more of the EIP completing in the average mosquito lifespan. Daily values in each scenario were smoothed with a seven-day rolling average to prevent stochastic hotspots from inflating risk. Values for each model run were averaged to give a yearly projected estimate under RCP8.5 [31].

## Results

To estimate the temperature-dependent USUV EIP, we modelled empirical infection data from a total of 565 *Cx. pipiens molestus* mosquitoes across 6 constant temperature treatments ranging from 17 to 25°C, from three previously published laboratory studies [5, 29, 30], as summarised in **Figure S1**. In these studies, adult mosquitoes were fed blood meals infected with USUV and maintained at these constant temperatures representative of UK climatic conditions. Subsets of mosquitoes were sampled at pre-determined intervals post blood meal and tested for viral RNA using RT-qPCR. We used data from one of the studies [30] to estimate the delay between initial body infection and systemic dissemination. This allowed us to modify the whole-body positivity data to estimate the temperature-dependent EIP of disseminated infection [5, 29, 30].

Model selection supported a model structure where the EIP_50_ varied log-linearly with temperature and the probability of infection was study-specific but did not vary with temperature. The looic results are shown in **Table 1**, where the lowest value indicates the preferred model structure. The parameters estimated by the model for this log linear function are given in **Table 2**, alongside the other parameters estimated by the model. It is worth reflecting that the model including both a log-linear median EIP *and* a logit-linear probability of infection scores similarly to the chosen model. It is intuitive that the infection probability *ρ* would vary across temperatures. However, given the number of temperature treatments and mosquitoes per treatment in the experiments, it is unsurprising that this extra parameter does not explain enough variation to merit inclusion in the model.

**Table 1:** Model selection results for Usutu EIP model using looic.

|  | Model structure | looic |
| --- | --- | --- |
| 1 | Approximate median EIP constant across temperatures. | 181.54 |
| 2 | **Approximate median EIP varies log-linearly with temperature.** | **148.56** |
| 3 | Approximate median EIP constant across temperatures, the probability of becoming infected varies logit-linearly with temperature. | 157.64 |
| 4 | Approximate median EIP varies log-linearly with temperature and the probability of becoming infected varies logit-linearly with temperature. | 150.23 |

**Table 2:** Parameters estimates for the Usutu EIP model.

| Parameter | Value<br>(posterior median with 95% credible intervals) |
| --- | --- |
| **Log-linear temperature dependent median EIP** |  |
| $\beta_0$ | 3.19 (95% CrI: 2.99 to 3.43) |
| $\beta_1$ | -0.55 (95% CrI: -0.8 to -0.36) |
| $\bar{T}$ | 21.03 |
| $sd(T)$ | 2.55 |
| **Intercept correction to reflect disseminated infections** |  |
| $\beta_2$ | 0.16 (95% CrI: -0.17 to 0.5) |
| **Weibull shape** |  |
| $\alpha$ | 1.44 (95% CrI: 0.97 to 2.11) |
| **Bloodmeal positivity** |  |
| $k$ | 0.24 (95% CrI: 0.19 to 0.31) |
| **Infection rates** |  |
| $\rho_{S=\text{Pilgrim}}$ | 95.67% (95% CrI: 77.44 to 99.69) |
| $\rho_{S=\text{Seechurn}}$ | 96.00% (95% CrI: 81.47 to 99.72) |
| $\rho_{S=\text{Holicki}}$ | 96.14% (95% CrI: 83.91 to 99.76) |
| $\bar{\rho}$ | 3.12 (95% CrI: 1.80 to 4.54) |
| $\sigma_\rho$ | 0.51 (95% CrI: 0.02 to 2.69) |
| **Beta-binomial overdispersion** |  |
| $\theta$ | 28.09 (95% CrI: 13.32 to 66.53) |

In **Figure 1** we show the model fit and the inferred processes: the estimated model components (**A-C**), and how the structure fit to the qPCR positivity of the data (**D-E**). We estimate an exponential decline in the rate of positive tests from the bloodmeal, falling to 50% after around 2.5 days and becoming negligible towards two weeks post infection (**Figure 1A**). This detection rate includes both live virus that has not yet infected the mosquito and RNA that may not successfully infect the mosquito. In **Figure 1B**, we show how the cumulative probability of infection (described by the estimated Weibull probability density function) changes across the minimum and maximum experimental temperatures, with significantly steeper increases following infection observed across the relatively modest six-degree temperature interval used in the study. Combining these two functions (and the study specific probability of infection, ρ_*s*_), the model can describe the trends in body PCR positivity well, shown in **Figure 1E**. An initially high positivity following infection declines due to bloodmeal RNA being metabolised. Further along the infection timeline (particularly in hotter treatments), this decline is mitigated by an increase in the genuine positivity from infections. The model predicts a high probability of infection and low variance across study specific probabilities of infection (**Table 2**, all over 95%). This suggests that the experimental results from different authors were compatible. However, the model also explores lower infection rates, down to approximately 80%. This means that model posteriors reflect more incomplete infection rates in the credible intervals. **Figure 1D** shows the model fit to the leg / wing positivity data used to estimate the correction for disseminated infections.

We find that the extrinsic incubation period of Usutu virus becomes shorter at higher temperatures. The median EIP (EIP_50_), defined as the time after which 50% of mosquitoes are expected to be infectious, fell from 68.06 (95% CrI: 38.62 to 141.04) at 17 °C to 12.35 days (95% CrI: 8.10 to 17.09) at 25°C, as seen in **Figure 2**. We compared this estimated median function for the EIP_50_ of USUV to that of WNV [28], finding that USUV was 37.75% (95% CrI: 3.38 to 55.68) faster between the 17°C to 25°C temperature range. There was only 2% overlap in the posterior draws, indicating a high level of confidence that USUV has a faster EIP than WNV.

**Figure 2:**
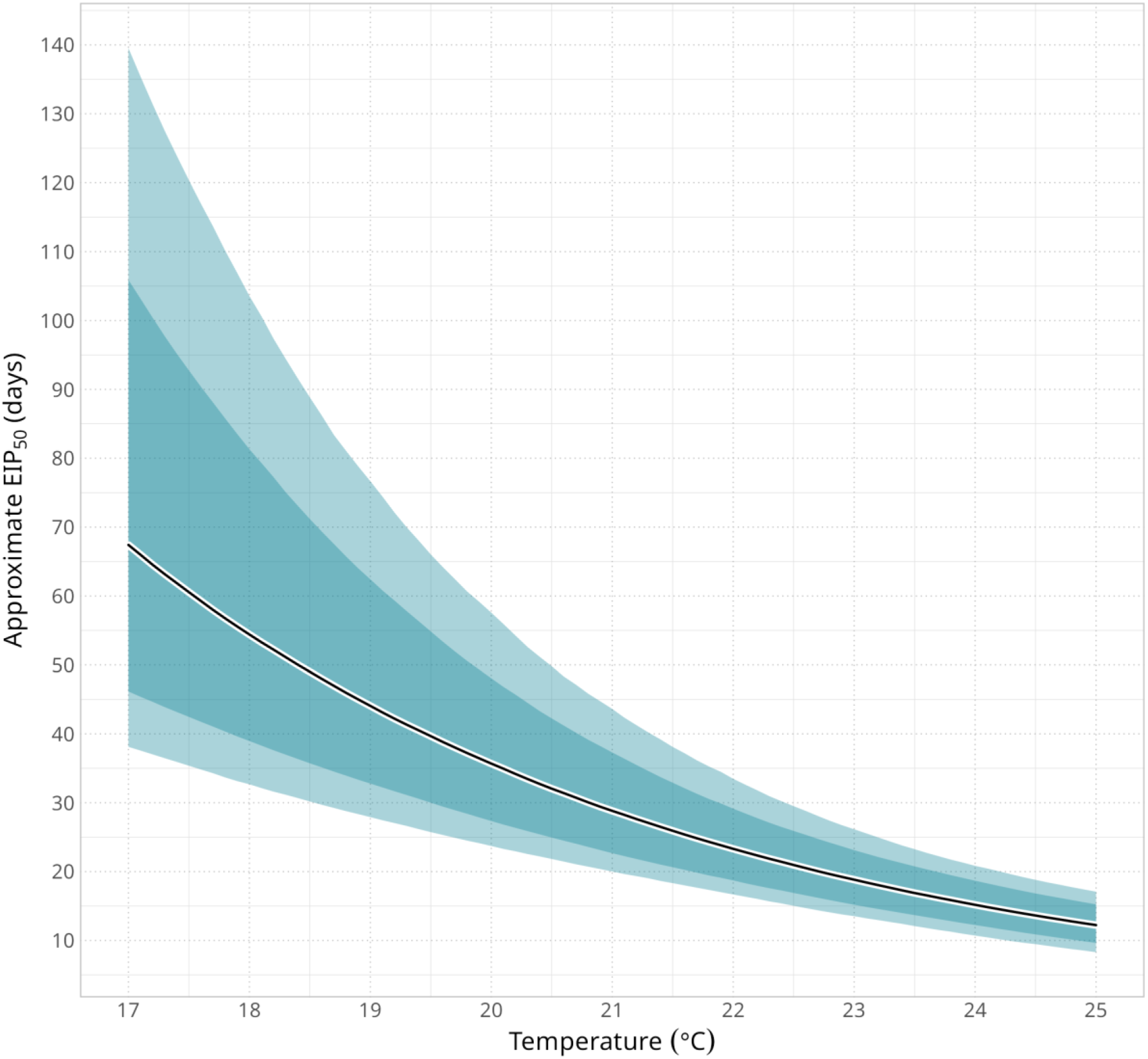
Temperature sensitive extrinsic incubation period of USUV. Modelled median approximate extrinsic incubation period (EIP_50_). The posterior median is shown as a line while 80% and 95% credible intervals are shown as dark and light shaded ribbons, respectively. Parameters for this function are given in **Table 2**.

We used our characterisation of the USUV EIP to compare the likelihood of USUV transmission and WNV transmission in the UK. Our findings suggest that in the temperate climate experienced in the UK, USUV transmission is more feasible than WNV transmission. For both viruses, we estimated a simple, data-driven metric: the probability that the EIP could complete within the average vector lifespan. We estimated the temperature-dependent survival in the mosquito biotype used in the empirical EIP studies (*Cx. pipiens molestus* for USUV and *Cx. pipiens pipiens* for WNV), using published experimental data [40, 41]. We combined these with the USUV EIP characterised in this study and our previously published estimates of the WNV EIP. It was regrettable that we were unable to compare the viruses in the same biotype of *Culex pipiens*, but we determined it was better to reflect the specific thermal biology of the biotype used in the corresponding laboratory studies rather than ignoring these differences altogether.

In **Figure 3** we compare the approximate USUV EIP in this paper with the values for WNV estimated by Vollans *et al.* (2024) [28]. We compare the pathogens across UK summer months (June to September), averaging across 2019 to 2024 temperatures. Then, we calculated the mean number of days in each month where it was more than 50% likely that the EIP would complete within the average vector lifespan. USUV exhibits a shorter monthly average EIP across the UK than WNV; the USUV EIP fell as low as approximately 45.97 days in parts of central and southeastern England in August, compared to a minimum of approximately 62.38 days for WNV. There were a greater number of days across the average summer season where the EIP of USUV could complete within the lifespan of *Cx. pipiens molestus* than for WNV in *Cx. pipiens pipiens*. For example, in August there were an estimated 22.33 days where there was a greater than 50% chance of the USUV EIP completing in the lifespan of *Cx. pipiens molestus*, compared to only 13.50 days for WNV in *Cx. pipiens pipiens*. Results for the survival models used to produce these plots are given in **Figure S2** and **Table S2**. In brief, the average lifespan of *Cx. pipiens molestus* fell from 75.80 days (95% CrI: 70.55 to 80.99) at 15°C to 18.39 days (95% CrI: 14.11 to 22.78) at 30°C. This compared to 93.36 days (95% CrI: 77.38 to 114.33) and 15.89 days (13.46 to 18.84) across the same temperature range for *Cx. pipiens pipiens*.

**Figure 3:**
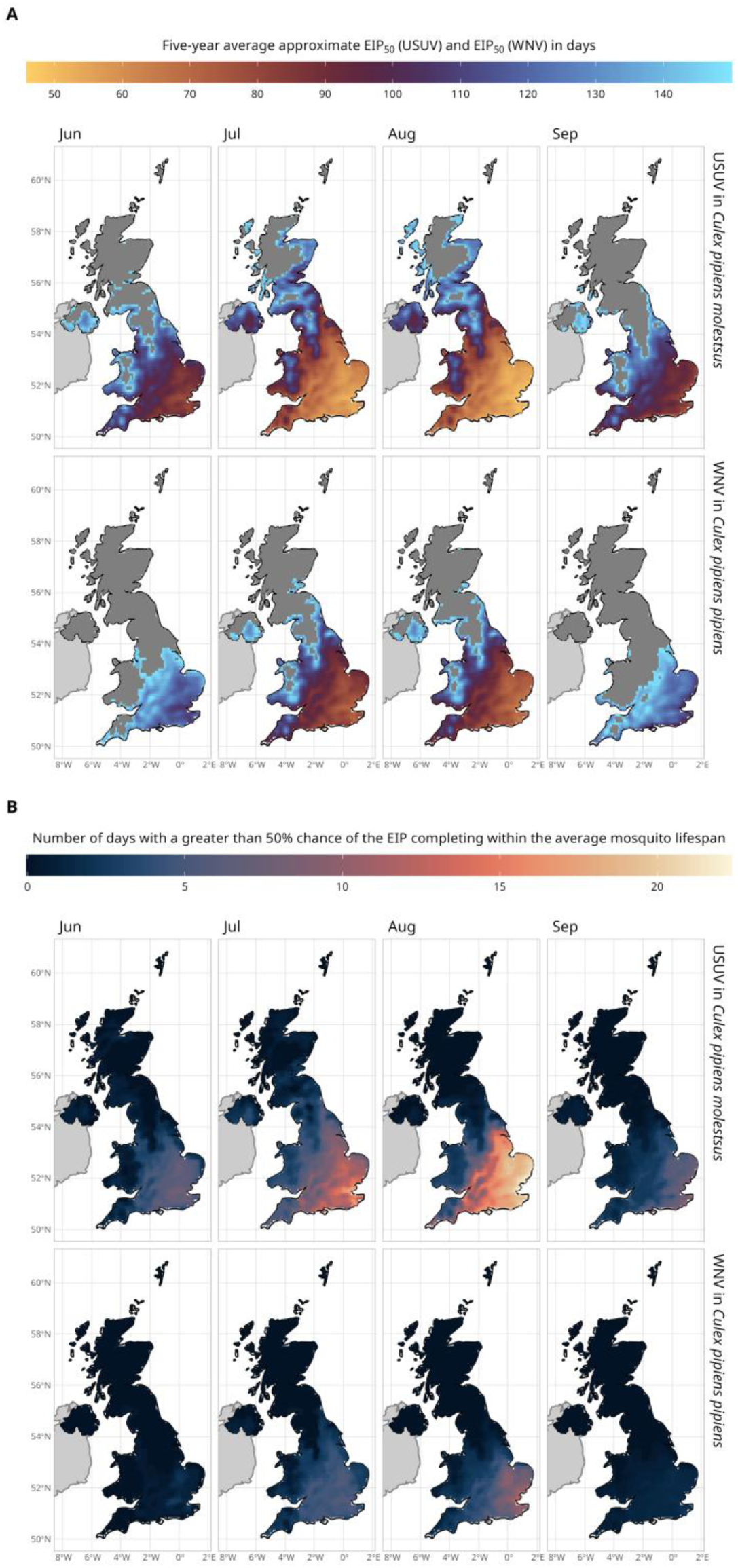
Seasonal variation in transmission risk. *Panel A* shows the monthly EIP_50_ of USUV (top row) and WNV (bottom row) in the UK. This is an inter-year average from 2019 to 2024. For both pathogens, these values were calculated using the log-linear function describing the EIP_50_ for each virus on daily temperature data. We calculated the monthly geometric mean across years from these daily values to give an average. Hotter colours indicate faster average EIP_50_ values and cooler colours slower EIP_50_ values. Grey regions show values above 150 days. *Panel B* shows the number of days where there was a greater than 50% chance that the estimated EIP could complete in the average mosquito lifespan for the temperatures in that month. Again, these values are the 2019 to 2024 average. Note that each pathogen is paired with the specific survival model for the species used in the corresponding experiments (*Cx. pipiens molestus* for USUV and *Cx. pipiens pipiens* for WNV). Hotter colours indicate more days above the 50% probability threshold.

In **Figure 4** we summed the number of days in the year where there was a greater than 50% chance of the EIP completing within the vector lifespan. Averaging across 2019-2024 there were 59 days where the USUV EIP could complete within the average lifespan of *Cx. pipiens molestus* compared to 24 days for WNV in *Cx. pipiens pipiens*. Approximately 37.34% of the UK landmass experienced at least 14 days where there was a 50% chance of the USUV EIP completing in the vector lifespan, compared to only 7.70% for WNV. For USUV, 23.29% of the country experienced 28 days’ worth of risk and 6.99% 42 days of risk, primarily concentrated in the south and east of England. Nowhere in the UK experienced either of these levels of risk for WNV.

**Figure 4:**
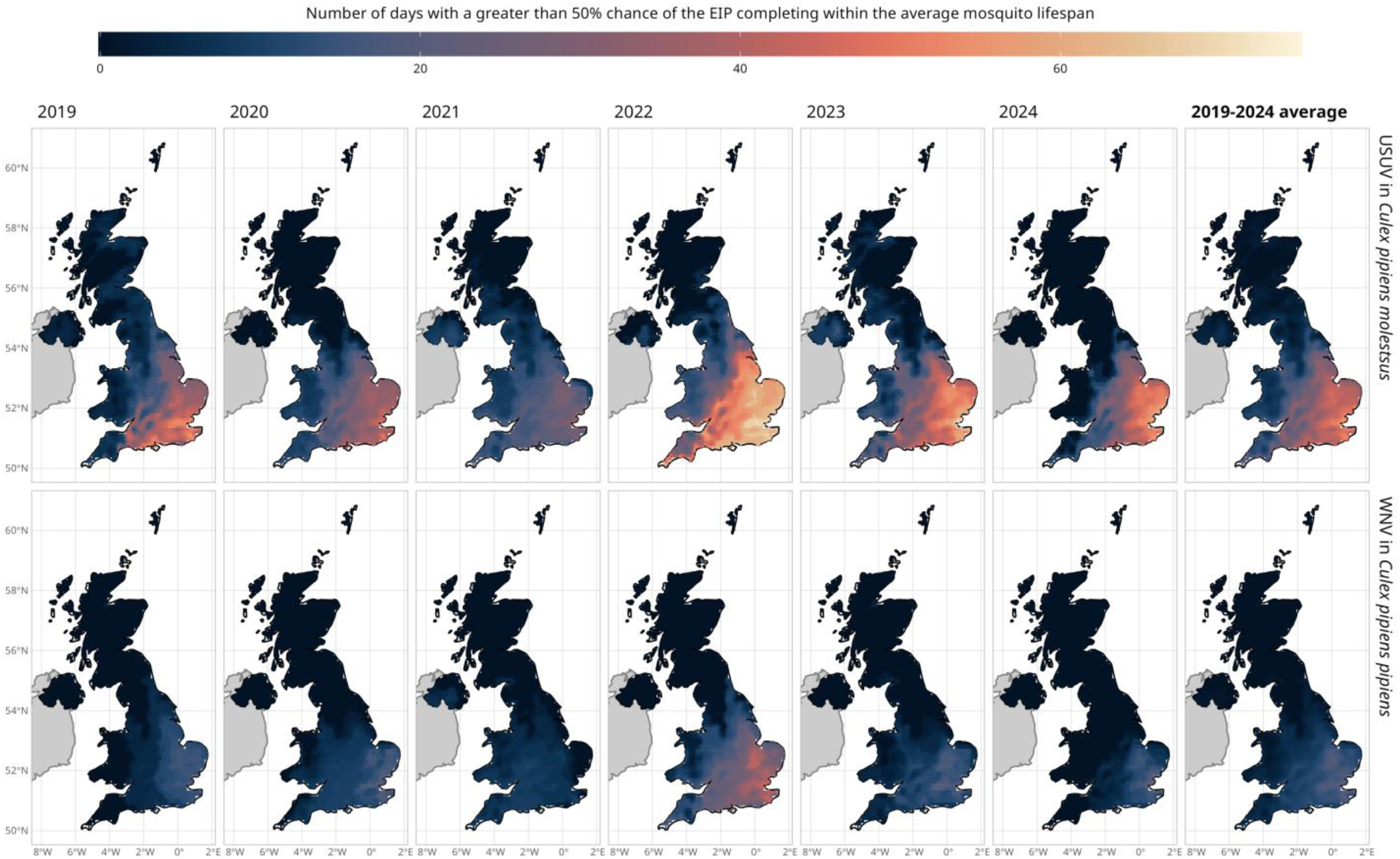
Annual variation in transmission risk. The colour temperature shows the number of days in each year (2019 to 2024) where there is a greater than 50% chance of virus EIP completing in the average mosquito lifespan of *Cx. pipiens molestus* (USUV, top row) and *Cx. pipiens pipiens* WNV, bottom row) in the UK. The final column shows the 2019 to 2024 average.

In **Figure 5 and 6** we use RCP8.5 climate projections for the UK to try to quantify how the transmission dynamics of USUV and WNV could change as temperatures increase under this specific emissions pathway. Values presented are averages of twelve probabilistic model runs. The number of days where there is a 50% chance or more of the EIP completing in the mosquito lifespan increases from a national average of 28.96 days in 2025 to 59.22 days in 2075 for USUV, compared to 13.99 days and 35.20 for WNV for the same years. The projected scenario suggests that by 2075, 72.58% of the UK will experience at least 28 days of USUV transmission risk, expanding into Scotland, Wales and Northern Ireland. The same statistic is 50.35% for WNV, an expansion which is a doubling of the 24.17% share of the country in 2025. **Figure 6** differences the number of days transmission risk for USUV in 2025 from the number of days for WNV in each projected year; with Usutu currently endemic, this highlights when a similar level of endemic transmission risk could be reached. These results suggest that between 2055 and 2065 WNV will experience a similar number of days at the 50% risk threshold as USUV in 2025.

**Figure 5:**
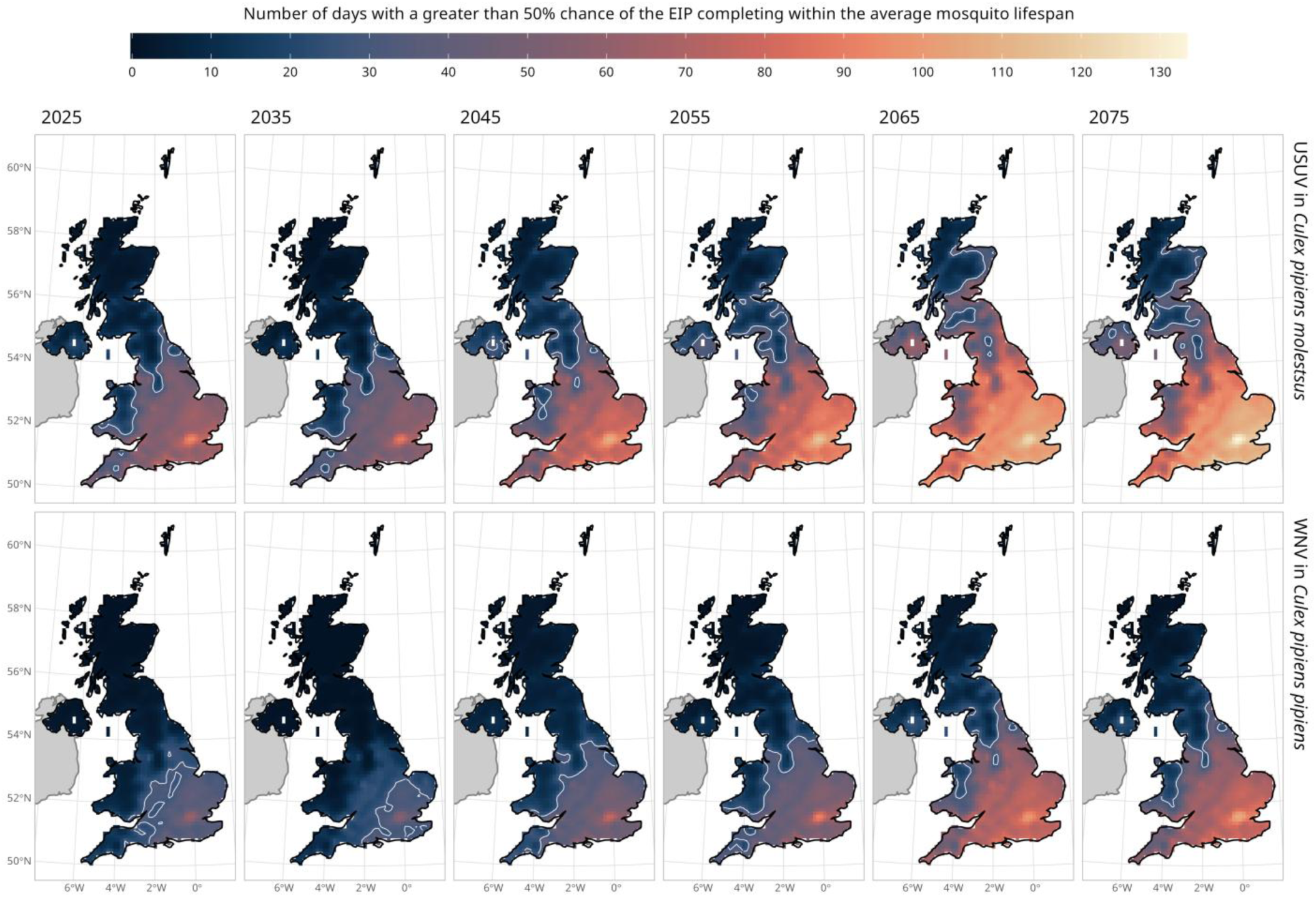
Expanding and increasing transmission risk under climate change. The maps show the number of days in each year where there is a greater than 50% chance of the EIP of the virus completing in the average mosquito lifespan of *Cx. pipiens molestus* (USUV, top row) and *Cx. pipiens pipiens* (WNV, bottom row) in the UK. Each column shows a decade from 2025 up until 2075 for RCP8.5. Values are the average of twelve probabilistic runs for each given year. There is stochastic variation in individual years that causes particular years to deviate from the general trend show across decadal increments. A white contour line delineates the region where there is a least a month (28 days) of the year where the 50% probability condition is expected to be met. Note the small area of white in Northern Ireland is Lough Neagh.

**Figure 6:**
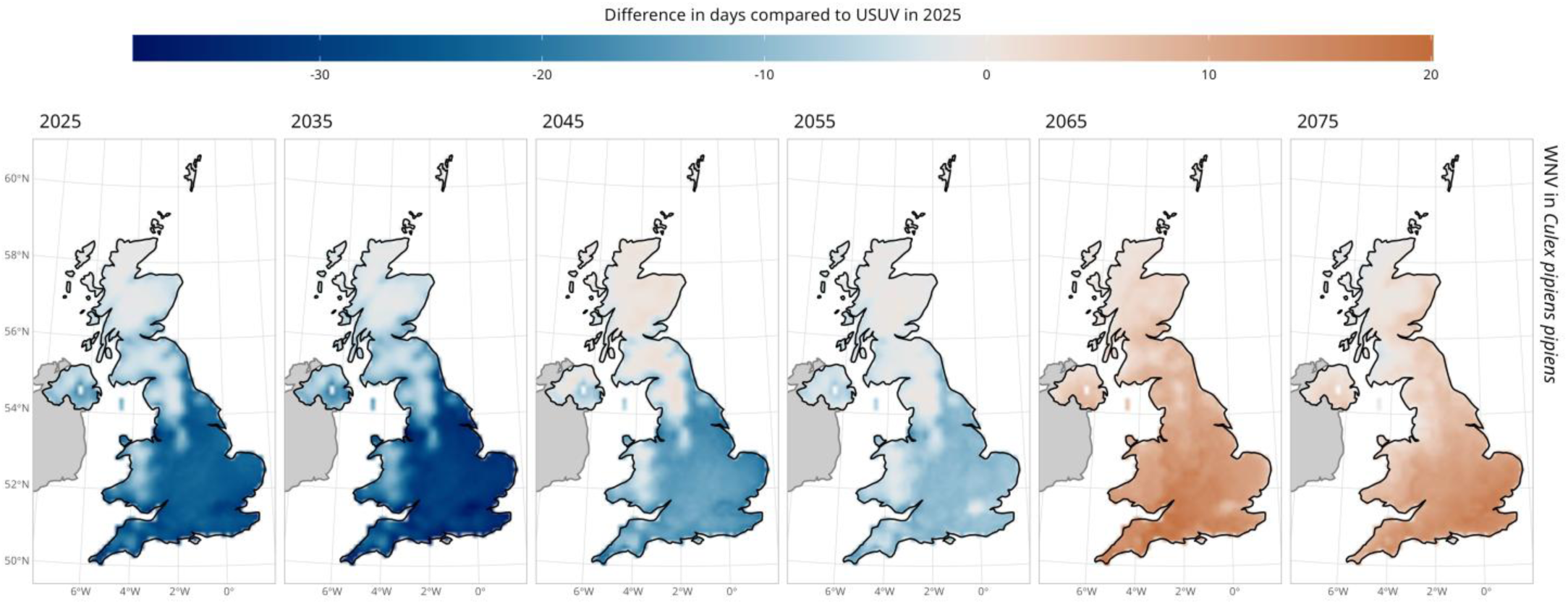
Evolving timeline of WNV transmission risk between 2025 and 2075, compared to USUV risk in 2025. The maps show the difference between the transmission risk of WNV compared to USUV in 2025, where blue indicates ower risk, white equivalent risk and red a higher risk. We calculated the number of days in 2025 where the EIP of USUV had a 50% or greater chance of completing in the mosquito lifespan for each grid cell. This was then subtracted from he equivalent value for WNV in each year to give the number of days less or more than the current USUV transmission isk. Note the small area of white in Northern Ireland is Lough Neagh.

## Discussion

Usutu virus (USUV) is an emerging mosquito-borne virus of growing public health concern, with a geographic range that has expanded northwards across Europe in recent years. Despite this, key epidemiological parameters governing USUV transmission remain poorly characterised. Consequently, modelling of USUV has relied on surrogate parameters from other members of the Japanese encephalitis virus complex, commonly West Nile virus (WNV) [42, 43]. This introduces substantial uncertainty, limiting our ability to accurately capture USUV-specific transmission dynamics. Across Northern Europe, USUV transmission (often alluded to by die offs in avian hosts) precedes the detection of WNV, making it a priority that we understand why this is the case. Here, we address this knowledge gap by providing the first characterisation of the temperature-dependent extrinsic incubation period (EIP) of USUV in the bridge vector *Culex pipiens molestus*. Using this model, we compared the feasibility of USUV and WNV transmission in the UK, potentially elucidating why USUV is currently endemic while WNV has not yet known to have established.

Across the temperature range examined (17 to 25°C), we find that the median USUV EIP decreases log-linearly with temperature, more than halving from 68.06 days (95% CrI: 38.62 to 141.04) at 17°C to 12.35 days (95% CrI: 8.10 to 17.09) at 25°C. The strong thermal sensitivity of the USUV EIP is expected: mosquitoes are poikilotherms, and their internal temperature varies with ambient temperature. Consequently, higher environmental temperatures can accelerate viral replication and dissemination, leading to shorter EIPs. Characterising temperature-dependent parameters for vector-borne pathogens is essential for assessing transmission risk: the EIP is a key metric underpinning measures such as vectorial capacity and the basic reproduction number (R₀) [44].

We used our characterisation of the USUV EIP to compare the likelihood of USUV transmission and WNV transmission in the UK. Our findings suggest that in the temperate climate experienced in the UK, USUV transmission is more feasible than WNV transmission. For both viruses, we estimated a simple, data-driven metric: the probability that the EIP could complete within the average vector lifespan. We did this by characterising temperature-dependent survival in the mosquito biotype used in the empirical EIP studies (*Cx. pipiens molestus* for USUV and *Cx. pipiens pipiens* for WNV) and combining these with the USUV EIP characterised in this study and previously published estimates of the WNV EIP [28]. It was regrettable that we were unable to compare the viruses in the same *Culex pipiens* biotype, but we determined it was better to reflect the specific thermal biology of the biotype used in the corresponding laboratory studies rather than ignoring these differences altogether.

We concluded that the median EIP of USUV was 37.75% (95% CrI: 3.38 to 55.68) shorter than that of WNV, at least across the 17 to 25°C temperature conditions explored here. This supports the observation that USUV tends to establish earlier than WNV in temperate regions of Europe [21, 45]. Although both viruses circulate primarily in enzootic cycles between birds and *Culex* mosquitoes, USUV has been more successful in cooler climates such as the Netherlands [45], Denmark [46], Germany [47] and the UK [10, 12]. Until now, there has been no clear quantitative explanation for these differences. Our results suggest that the success of USUV in cooler regions may, in part, be explained by a faster EIP in temperature climates, allowing the virus to infect and disseminate within the lifespan of the vector. Indeed, the outcome of WNV strain competition in the northern United States was determined by an emergent strain replicating faster at colder temperatures [28, 32]. This suggests that faster replication is a valuable attribute in more temperate climates.

We then examined the risk of transmission of WNV and USUV under RCP 8.5 climate projections. Our results suggest a progressive geographic expansion in the region of thermal suitability for both pathogens from 2025 until 2075. We show that by between 2055 and 2065 WNV could meet or exceed the number of days where the EIP is likely to complete in the mosquito lifespan compared to USUV in 2025. We must emphasize that meeting or exceeding the risk we present for USUV in 2025 does not guarantee or preclude the establishment of WNV in the UK; it is simply an indicative benchmark that if another mosquito borne pathogen (USUV) can become endemic under these conditions, then it is credible that another that meets these conditions also could.

Our USUV EIP model is the first to provide a robust method for estimating the EIP from qPCR body-positivity data. Our model primarily relies on temperature-dependent whole body positivity data measured by RT-qPCR [5, 21], supplemented by a single-temperature study reporting body and leg / wing positivity [30]. While plaque assays are the gold standard for assessing mosquito infectivity, qPCR offers practical advantages: it is faster, cheaper, more scalable, and requires lower biosafety containment [48, 49], and has been used to study EIP in other arboviruses such as Zika virus [50]. While saliva or leg / wing samples would better reflect viral dissemination, saliva extraction is technically challenging [25], and temperature-dependent leg / wing positivity data is not available for USUV. This framework demonstrates how body positivity data can be leveraged by modelling blood-meal RNA decay to approximate true infection and applying a modifier to account for the delay between body infection and dissemination.

Our analysis relied on empirical data from three laboratory studies with experimental designs that required specific statistical treatment. This meant we have had to make a series of compromises and assumptions. First, this model parameterisation relied on data collected across a relatively narrow temperature range, highlighting the necessity for further empirical work to validate applicability across broader temperature regimes, including future UK climates and other countries. Second, the USUV EIP data were collected in a “bridge vector”, *Cx. pipiens molestus.* This biotype is interesting as it is important for human spillover risk, with diverse biting habits including both birds and mammals [51]. However, as vector competence can vary by biotype and species [5], it would be useful to collect high quality empirical data on USUV EIP in *Cx. pipiens pipiens*, the primary biotype maintaining USUV in the avian reservoir. Third, our model relies heavily on body positivity data, and the modifier used to approximate dissemination is based on a single temperature treatment. Temperature-dependent data on EIP proxies such as leg dissemination, in our view, remain a more appealing way of measuring EIP than either saliva or body positivity [28] and this remains an important avenue for further empirical work. Fourth, the empirical data used to characterise the WNV EIP and USUV EIP models vary. The WNV EIP model is based on data from plaque assays of mosquito legs [28, 32], while the USUV EIP model uses predominantly body positivity data from RT-qPCR, supplemented by leg positivity data. Additionally, the WNV EIP is characterised in colonies of *Cx. pipiens pipiens*, but the USUV EIP is estimated in colonies of *Cx. pipiens molestus*. Finally, the lab studies use different temperature treatments, rearing regimes, mosquito strains and USUV lineages. While there are terms in the model to try and account for inter-study variability (the study-specific probability of infection), this may not fully capture the extent of the underlying experimental variation. These differences have motivated us to be clear about the limitations of our approach and the methodological differences that may complicate comparisons across pathogens and vectors.

We explored the impact of climate change on the transmission dynamics of USUV and WNV, however we are limited by the available climate scenarios available. Our modelling metrics required a daily timestep and high spatial resolution, and the highest temporal and spatial resolution we could obtain was for Representative Concentration Pathway (RCP) 8.5. RCP 8.5 represents a trajectory in which emissions continue largely unmitigated. This ties our projections to a pessimistic future for USUV and WNV transmission risk based on this high-emissions scenario. In addition, our projections assume no evolutionary or ecological adaptation of either the mosquito vectors or the viruses. The extrinsic incubation period (EIP) and mosquito survival models are parameterised using data from contemporary populations and are applied directly to projected temperature changes. As a result, potential future changes in vector or viral thermal responses are not captured.

All coarse-scale analyses of arbovirus infection and ecology must approximate local dynamics. To estimate the EIPs and adult mosquito lifespans across the UK, and through time, we use the ambient air temperature. This implicitly assumes that mosquitoes experience these predicted air temperatures directly, and that this temperature determines viral dissemination and adult mosquito survival. This assumption may be violated by mosquitoes, which may moderate their experienced temperature across heterogeneous microhabitats. For instance, the *Cx. pipiens molestus* biotype commonly inhabits sheltered, belowground human-made structures such as subways, basements and sewage infrastructure year-round. Consequently, *Cx. pipiens molestus* may experience warmer and more thermally stable conditions than those captured by ambient air temperature, especially during cooler periods. Our temperature-based estimates may therefore underestimate transmission potential in some settings, through ignorance of experienced temperature of the pathogen in a vector routinely seeking shelter in locations with warmer temperatures. Furthermore, our analysis does not account for spatial variation in mosquito or avian host abundance. Because the primary transmission cycles of both USUV and WNV occur between mosquitoes and birds, realised transmission risk depends not only on temperature suitability but also on local host and vector densities. This is particularly the case for the establishment of WNV and reintroductions of USUV, where infected migratory birds from the continent can trigger cycles of infection. Our results should therefore be interpreted as characterising the climatic feasibility of transmission, rather than predicting the precise transmission intensity or vector habitat suitability.

## Conclusion

In this paper we approximate the temperature-dependent USUV EIP for the first time. This research contributes essential knowledge to a poorly characterised pathogen of increasing relevance to public health, particularly in Northern Europe. This USUV EIP can be used to estimate transmission risk and public health implications of the virus, with measures such as the basic reproduction number reliant on accurate pathogen-specific measures. Our evidence supports USUV as a precursor for WNV in Northern Europe and suggests that WNV endemicity will become increasingly likely with climate change, matching the dynamics of USUV by 2065 at the latest.

## Author contributions

RP, MV and JD conceived the study. RP designed and wrote the model with advice and review from MV, LG, MF and TW. LG, MV and RP extracted and interpreted the published data. MV and JD were project administration. RP, MV and LG wrote the manuscript. All authors reviewed and contributed to the final manuscript.

## Supplementary materials

### Survival model

We model observed lifespans, *L*, with a Weibull distribution where the inverse scale parameter (^1^⁄_*ω*_) is a log-linear function of standardised temperature and *φ* is the shape parameter:

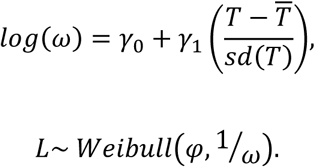

The intercept and gradient of the function are given by *γ*_0_ and *γ*_1_ respectively. We assume that the same shape parameter can be used to describe *Cx. pipiens molestus* survival times as was used to describe *Cx. pipiens pipiens* survival times but allow the temperature dependent function to vary across the two biotypes. As we only had mean and standard errors for each *Cx. pipiens molestus* temperature treatment, we implemented these data as a soft constraint prior on the predicted mean (*E*(*L̅*)) of the Weibull distribution:

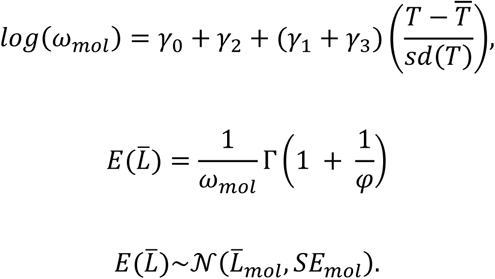

Here, *γ*_2_ and *γ*_3_ are the intercept and gradient adjustment for the temperature dependent response for *Cx. pipiens molestus.* The inputs *L̅*_*mol*_ and *SE*_*mol*_ are the mean and standard error from each experimental treatment. Priors for this model are given in **Table S2**.

**Table S1:**
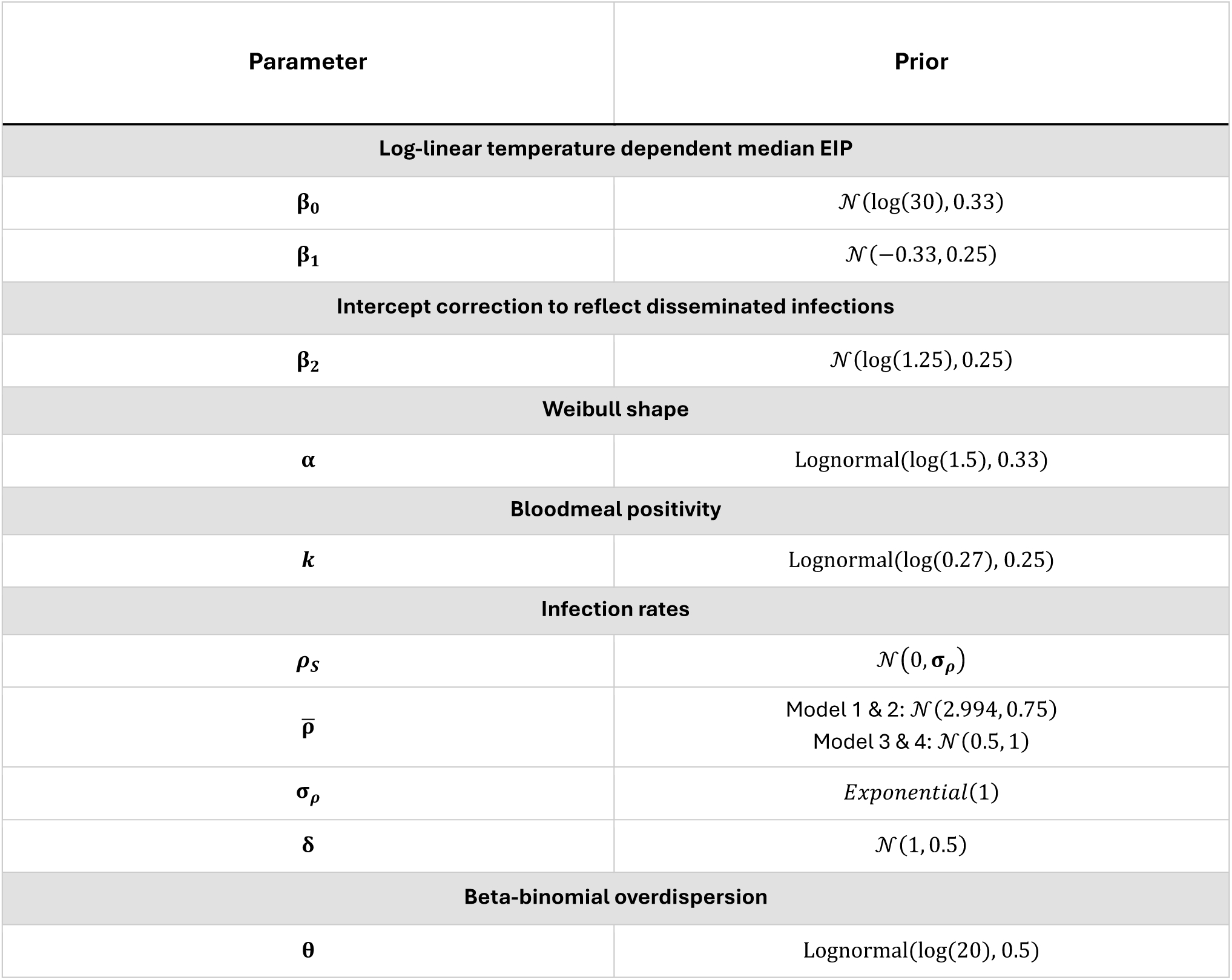
Priors used for the Usutu EIP model.

**Table S2:**
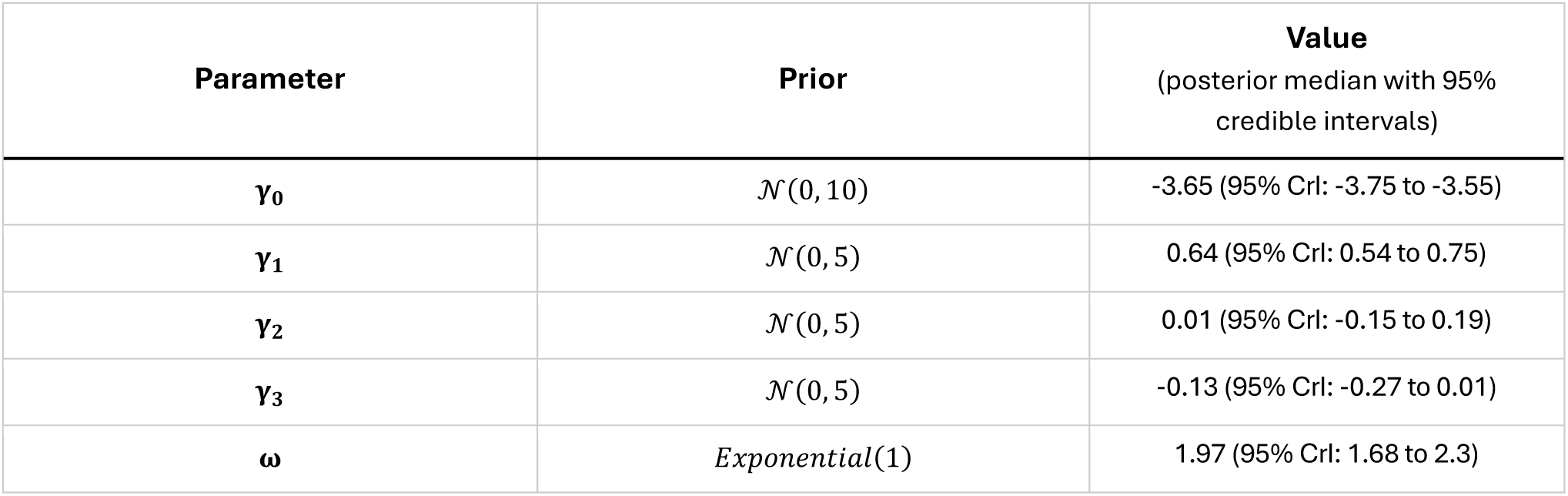
Priors used for the *Culex pipiens pipiens* and *Culex pipiens molestus* survival models.

**Figure S1:**
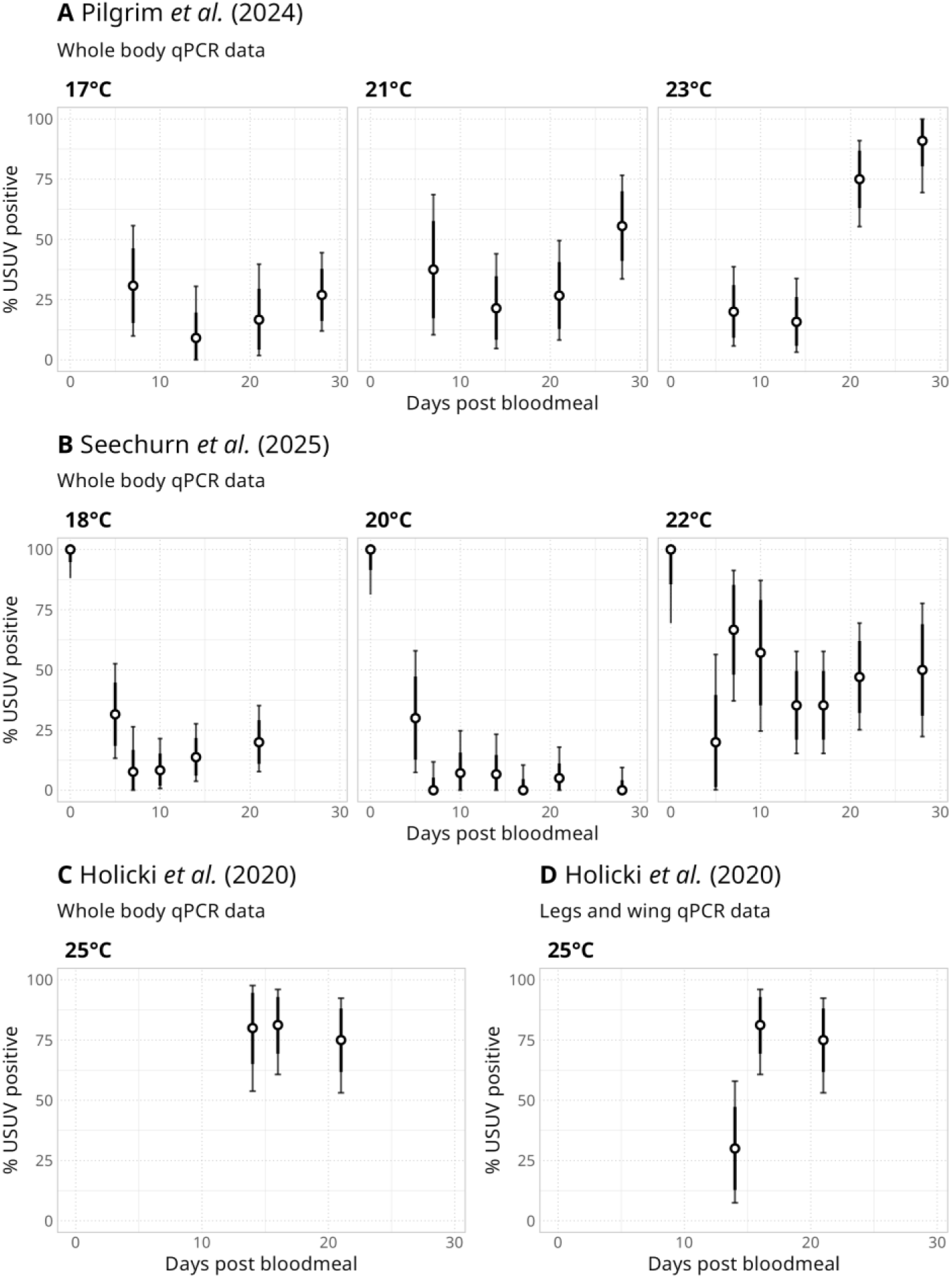
Data overview. An overview of the experimental data discussed in this study. Points indicate proportions calculated for each sample and thick / thin lines denote the binomially distributed 80 / 95% credible intervals for each proportion. **Panel A** shows body data from Pilgrim *et al.* (2024) and **panel B** Seechurn *et al.* (2025) (both sets of results refer to *Culex pipiens molestus*). Columns show different temperature treatments (note that temperature conditions are slightly different for each study). In **panel C** we show results for Holicki *et al.* (2020) for body positive PCR tests while **panel D** shows leg / wing positivity.

**Figure S2:**
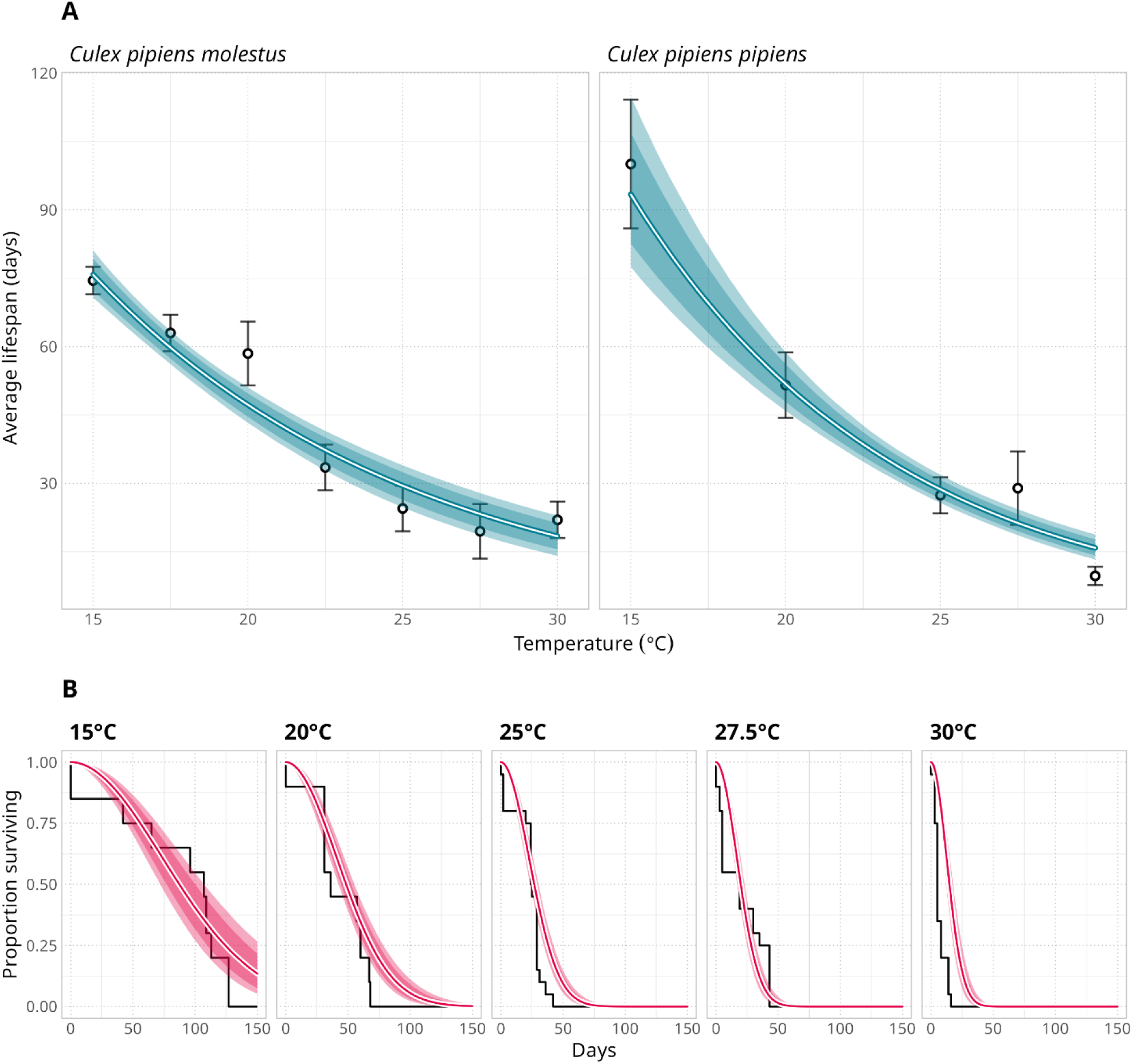
We estimated the temperature dependent survival functions for *Cx. pipiens pipiens* and *Cx. pipiens molestus.* We digitised row-level data for the lifespan of *Cx. pipiens pipiens* from Andreadis *et al. (*2014) and used priors from plots in Spanoudis *et al.* (2018) to adjust the model for *Culex pipiens molestus*. In Panel A we show the temperature dependent mean lifespan of each vector. Points and intervals show the priors used for *Cx. pipiens molestus* (mean lifespan as points and standard error as bars) while these points and intervals reflect summaries of the pseudo row level data extracted for *Cx. pipiens pipiens*. In *Panel B* we show the model fit for the pseudo row level data for *Cx. pipiens pipiens*.

